# Comprehensive analysis of HIV elite controllers’ and progressors’ transcriptional profiles from CD8^+^ T lymphocytes demonstrates heterogeneity of pathways and master regulators, which may be essential for disease nonprogression

**DOI:** 10.1101/2020.10.19.346528

**Authors:** Sergey Ivanov, Dmitry Filimonov, Olga Tarasova

**Author notes:** Address correspondence to Sergey Ivanov.

## Abstract

Cytotoxic and noncytotoxic CD8^+^ T lymphocyte responses are essential for the control of HIV infection. Understanding the mechanisms underlying HIV control in elite controllers (ECs), who do not progress to AIDS for many years without treatment, may facilitate the development of new effective therapeutic strategies.

We performed a comprehensive analysis of the transcriptional profiles of CD8^+^ cells from ECs and treated and untreated progressors using an original pipeline. Cluster analysis enabled the identification of five distinct groups (EC groups 1-5) of ECs based on their transcription profiles. Profiles of EC groups 2-4 were associated with different numbers of differentially expressed genes, but the corresponding genes shared the same cellular processes. These three groups were associated with increased metabolism, survival, proliferation, and the absence of an “exhausted” phenotype of CD8^+^ lymphocytes, compared to untreated progressors. The transcriptional profiles of EC group 1 were opposite to those of EC groups 2-4 and similar to those of the treated progressors. This group may be associated with residual dysfunction of T lymphocytes. The EC group 5 was indistinguishable from normal. EC groups 1 and 5 can have mechanisms of nonprogression unrelated to CD8^+^ cells.

The transcription changes in CD8^+^ lymphocytes from ECs may be attributable to the receptors that modulate the functional states of CD8^+^ cells. We identified 22 receptors, which are potential master regulators and may be responsible for the observed expression changes of CD8^+^ cells in ECs. These receptors can be considered potential targets of therapeutic intervention, which may decelerate disease progression.

**IMPORTANCE:** A small group of HIV-infected individuals, known as elite controllers (ECs), do not develop AIDS for many years, despite being untreated. Understanding the mechanisms governing HIV control in ECs may help develop new effective therapeutic strategies. Since CD8^+^ T lymphocytes seem to play a significant role in HIV control, we analyzed the large number of transcriptional profiles in total CD8^+^ cells from ECs, treated, and untreated progressors. We found distinct groups of ECs, which may have different mechanisms governing HIV nonprogression, and we identified related pathways and cellular processes that are dissimilar from those in progressors. We also identified master regulators, key proteins in the signaling network that can be responsible for observed transcriptional changes in CD8^+^ cells from ECs and may be essential for disease nonprogression. These proteins may represent potential targets for therapeutic interventions.

## INTRODUCTION

Human immunodeficiency virus (HIV) infection remains one of the most significant challenges facing humankind. Approximately 38 million people were living with HIV, and 690 thousand died by the end of 2019 (https://www.who.int/news-room/fact-sheets/detail/hiv-aids). Early administration of combined antiretroviral therapy (cART) enables HIV infection to switch to a chronic form and increases the lifespan of patients to those seen in uninfected people; however, irregular use of cART and adverse effects caused by particular drugs and their toxicity significantly hinder the efficiency of this approach (1–3). Moreover, since the presence of latent HIV infection, cART must be administered throughout life because interruption of therapy leads to viral rebound and disease progression (4, 5). Thus, new approaches to treat HIV infection are being developed (5–9). One of the most promising approaches is developing therapeutic and preventive vaccines; however, no effective vaccines exist at present (6). This lack of a vaccine can be explained by the fact that people do not develop natural, protective immunity to HIV infection, whereas almost all successful vaccines were created for diseases for which natural immunity exists (8). Since some vaccine candidates allow moderate protection from HIV, a successful vaccine may potentially be developed; however, to this end, a more profound understanding of the interaction between HIV and the immune system is required.

HIV infects CD4^+^ T helper cells, as well as monocytes and macrophages and various kinds of dendritic and epithelial cells (10). HIV infection causes innate and later adaptive immune responses, where the latter includes both cellular and humoral components. The formation of anti-HIV antibodies may be essential for control under viremia. On the other hand, the CD8^+^ cellular response seems to have a significant role (11). The cellular response causes a decline in viral load in the acute phase of infection but cannot eliminate the virus from the organism because a high mutation rate during replication allows HIV to escape from the immune response. Persistent infection causes chronic activation of HIV-specific CD4^+^ and CD8^+^ cells, which leads to their apoptosis (12, 13). A phenomenon called “bystander activation” was described in chronically infected patients, in which significant numbers of CD4^+^ and CD8^+^ non-HIV-specific T lymphocytes are activated in a T-cell receptor-independent and cytokine-dependent manner that leads to their apoptosis (11, 14–16). Chronic infection also causes the dysfunction or “exhaustion” of CD8^+^ lymphocytes, which is characterized by increased expression of inhibitory immune checkpoints PD-1, Lag-3, and Tim-3 on the surface of cells and decreased ability of CD8^+^ cells to proliferate, secrete cytokines, and induce cytotoxicity (17). The high mutation rate of HIV, immune exhaustion and the loss of CD4^+^ and CD8^+^ cells make further immune responses ineffective with subsequent progression to AIDS (18). However, the time from infection to the development of AIDS can vary significantly among patients. Most infected patients, called progressors, usually develop AIDS after 8-10 years, but a small group of people, known as long-term nonprogressors (LTNPs), remains asymptomatic for more than ten years and are characterized by higher CD4^+^ cell counts and lower viral load (13, 19–21). LTNPs can be divided into several groups. The group, called elite controllers (ECs), demonstrates the best control of viral replication with a viral load less than 50 copies/ml while maintaining CD4 cell counts from 200 to 1000/ml. Little is known about the mechanisms of viral control in ECs (11, 13, 18–24), but it seems that host and viral factors, as well as various cell types, may be involved in control, and the cellular response of CD8^+^ lymphocytes has a major role.

Investigation of CD8-related mechanisms of viral control in ECs is essential because mimicking similar responses in chronically infected progressors may lead to the functional remission of HIV infection (25).

To date, most of the studies related to the investigation of mechanisms of HIV suppression in ECs have focused on particular molecules or pathways, whereas analysis of HIV-related genome-wide OMIC data provides an opportunity to reveal novel mechanisms that were not known or previously hypothesized (26). Transcriptomics studies are the most frequent in HIV research and include those investigating CD8^+^ lymphocytes from ECs (27–30). Most of the corresponding studies were focused on total CD8^+^ lymphocytes, rather than HIV-specific lymphocytes (27, 29, 30). This focus is important because non-HIV-specific CD8^+^ and CD4^+^ lymphocytes are involved in the pathogenesis of disorder by the “bystander activation” effect (see above), which leads to their dysfunction and apoptosis (11, 14–16). To identify specific HIV control mechanisms, the comparison of gene transcription in CD8^+^ lymphocytes from ECs to both progressors and healthy controls is required. For instance, Hyrcza and colleagues compared transcription profiles from total CD4^+^ and CD8^+^ lymphocytes between ECs, progressors (acute and chronic phases), and uninfected people (27). They found differentially expressed genes (DEGs) between ECs and progressors in both the acute and chronic phases but did not find differences between ECs and healthy controls. This can potentially be explained by the fact that CD8^+^ ECs have heterogeneous transcription profiles, and some of them are indistinguishable from healthy controls. Chowdhury and colleagues applied cluster analysis to CD8^+^ cell transcriptional profiles from 51 ECs and found five distinct groups (30). Some of the groups were distinguishable from both control samples and samples from cART-treated patients. The authors found that the pathways governed by mTOR and eIF2 proteins are potentially the most important for the functions of CD8^+^ lymphocytes in ECs and that these pathways are dominant in three out of five EC groups.

In the present study, we performed a comprehensive analysis of the transcription profiles of CD8 lymphocytes from a large cohort of ECs, cART-treated and untreated progressors using the pipeline, which includes the following steps: (1) identification of distinct groups of ECs based on the corresponding transcription profiles; (2) identification of differential expression for each EC group, cART-treated and untreated progressors compared to uninfected controls; (3) comparison of obtained DEGs between ECs and progressors; (4) identification and comparison of differentially regulated pathways in ECs and progressors; and (5) identification of master regulators (MRs), which are the proteins at the top of the signaling network regulating the expression of DEGs observed in ECs.

## RESULTS

### Identification of EC groups based on CD8^+^ cell transcription profiles

To identify distinct groups of ECs, we retrieved corresponding transcription data from CD8^+^ lymphocytes, as measured by Chowdhury and colleagues (30), from the Gene Expression Omnibus (GEO) (https://www.ncbi.nlm.nih.gov/geo). This dataset includes samples from 51 ECs, 32 cART-treated patients, and 10 uninfected individuals. We performed hierarchical clustering of 51 samples in the space of 7113 genes with the highest variance of expression values across samples (see Material and Methods) and found five distinct clusters (Fig 1) (see also Table S1). These clusters were generally similar to those obtained by Chowdhury and colleagues (30). Fig 1 shows that transcription profiles from ECs are heterogeneous, and some of the revealed groups have opposite profiles (“cyan” and “blue” groups).

**Fig 1.**
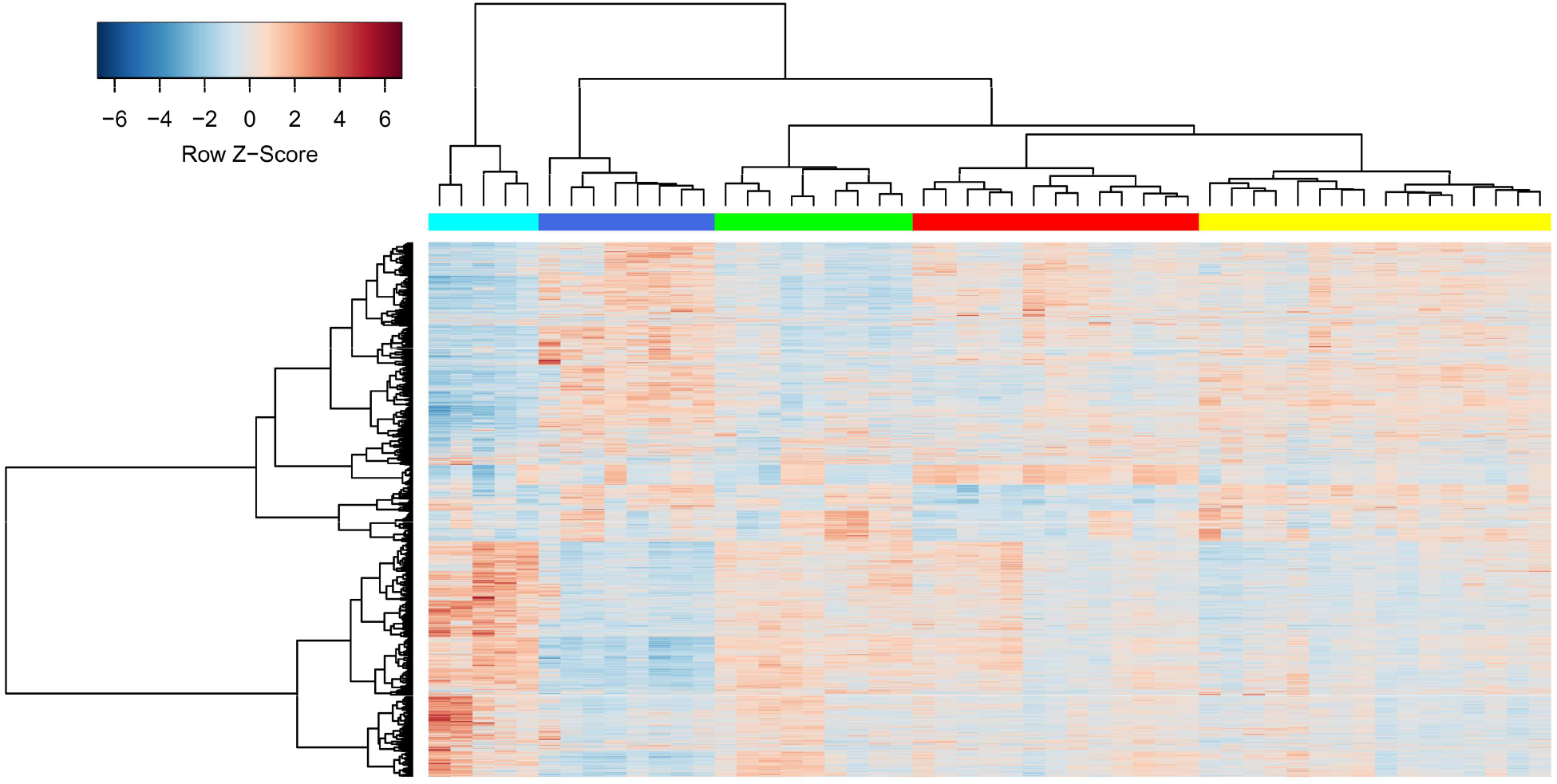
Heatmap demonstrating clustering results of CD8 transcription profiles from ECs. The rows in the heatmap are genes; the columns are samples. Row Z-Score is the number of standard deviations by which the value of gene expression in particular sample is above or below the mean value of all samples.

To estimate the significance of the obtained EC groups, we performed bootstrap resampling analysis and obtained p-values for each cluster: the higher the p-value, the more probable the existence of a cluster. Two types of p-values were calculated: the AU (Approximately Unbiased) p-value and BP (Bootstrap Probability) value, which were computed by multiscale and normal bootstrap resampling, respectively (see Materials and Methods). Most clusters have significant p-values of more than 0.95, and two clusters have p-values of more than 0.65 (Table 1), which is still acceptable for further analysis.

**Table 1.** The significance of EC clusters.

| Cluster No | Color in heatmap | AU p-value | BP p-value |
| --- | --- | --- | --- |
| 1 | Blue | 0.99 | 0.99 |
| 2 | Cyan | 0.66 | 0.65 |
| 3 | Green | 0.78 | 0.65 |
| 4 | Red | 0.99 | 0.98 |
| 5 | Yellow | 0.99 | 0.97 |

Hereafter, we will refer to EC groups by numbers in order as in Table 1 (EC groups 1-5).

### Identification of DEGs and their comparison between EC groups and cART-treated and untreated progressors

We identified DEGs between each of the EC groups and healthy controls as well as between cART-treated progressors and controls (Table 2). We also retrieved from GEO two other datasets with CD8^+^ cell transcription profiles from untreated progressors and corresponding healthy controls (see Materials and Methods). The first dataset (GEO ID: GSE6740), published by Hyrcza and colleagues, contains data on CD8^+^ lymphocytes from untreated progressors in acute and chronic phases of infection (27). The second dataset (GEO ID: GSE25669) contains corresponding information from untreated progressors in the acute phase. Since the transcription profiles were measured on different microarrays, we identified DEGs for each dataset separately. Numbers of up- and downregulated genes with log fold change > |0.7| and adjusted P-value < 0.1 in various groups of ECs and progressors are given in Table 2. The corresponding thresholds were chosen empirically to balance the number of DEGs and statistical significance of differential expression (for details, please see the Materials and Methods section). Only EC groups 2 and 3, as well as untreated progressors, were associated with a high number of DEGs, whereas other groups containing the most samples were only slightly different from healthy controls.

**Table 2.** Numbers of up- and down-regulated genes identified in groups 1-5 of ECs and progressors.

| Groups | No of samples | Up-regulated | Down-regulated |
| --- | --- | --- | --- |
| EC group 1 (blue) | 8 | 4 | 65 |
| EC group 2 (cyan) | 5 | 1063 | 1073 |
| EC group 3 (green) | 9 | 263 | 351 |
| EC group 4 (red) | 13 | 79 | 16 |
| EC group 5 (yellow) | 16 | 14 | 17 |
| cART-treated progressors | 32 | 14 | 86 |
| Untreated progressors (acute phase) 1 | 5 | 362 | 303 |
| Untreated progressors (acute phase) 2 | 4 | 331 | 479 |
| Untreated progressors (chronic phase) | 5 | 95 | 118 |

To compare transcriptional profiles between EC groups together with cART-treated progressors and healthy controls, we performed cluster analysis in the space of genes that were differentially expressed in at least one EC group or cART group compared to the healthy control (Fig 2).

**Fig 2.**
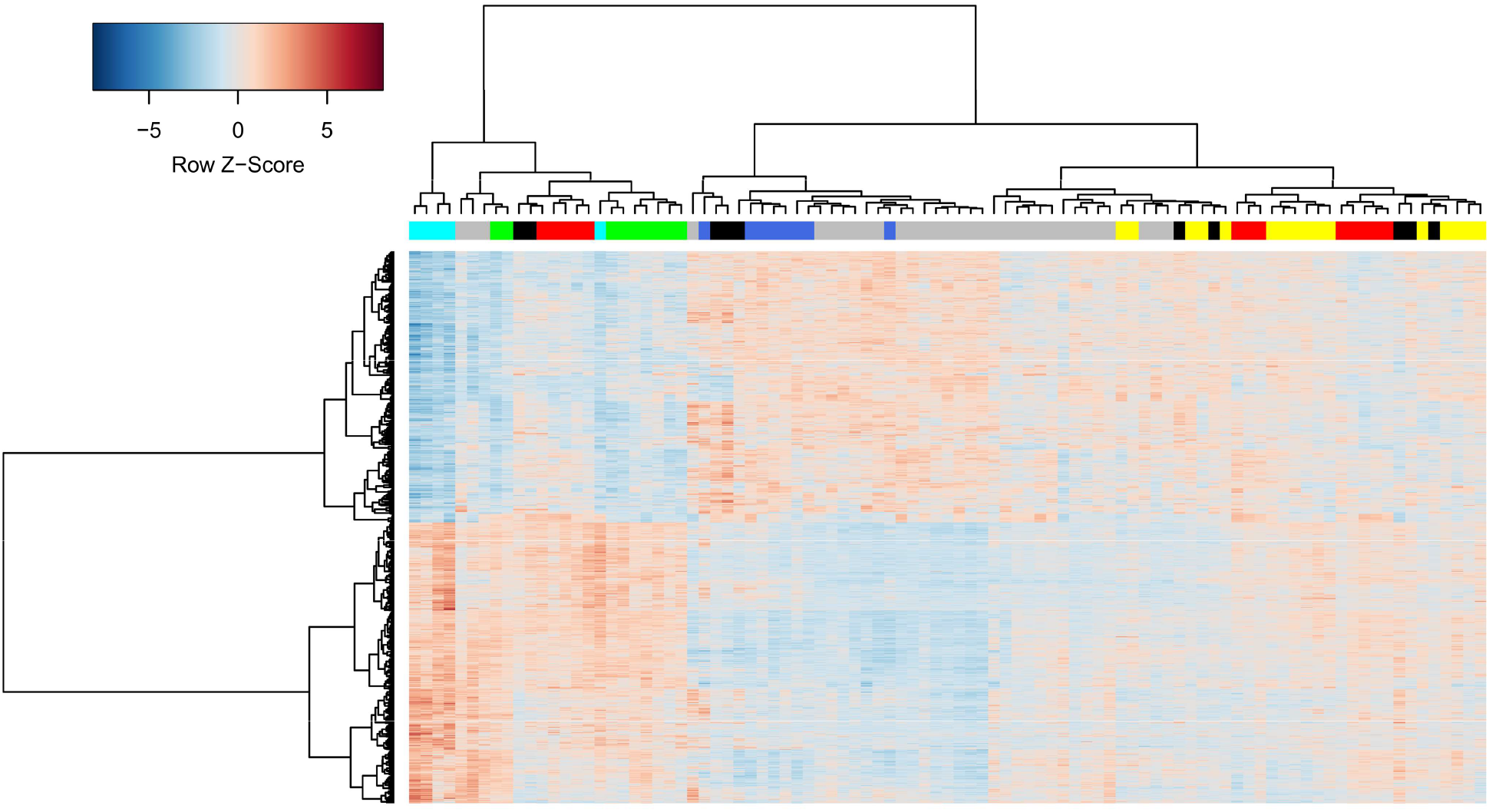
Comparison of five EC groups with cART-treated progressors and healthy controls. The rows in the heatmap are genes; the columns are samples. The blue, cyan, green, red, and yellow colors of columns represent EC groups 1-5 (Table 1); grey and black colors represent cART-treated progressors and healthy controls, respectively. Row Z-Score is the number of standard deviations by which the value of gene expression in particular sample is above or below the mean value of all samples.

Analysis of the content of Table 2 and Fig 2 allows the following conclusions to be drawn. First, all transcription profiles from EC group 5 and some profiles from EC group 4, as well as cART-treated progressors, are indistinguishable from healthy controls (right part of Fig 2). This finding means that total CD8^+^ lymphocytes from corresponding ECs are potentially not involved in control under viremia. On the other hand, considerable control under viral replication was achieved in corresponding cART-treated progressors. Since samples from cART-treated progressors are divided into two groups on the dendrogram (Fig 2), we performed bootstrap resampling analysis in a similar manner to the case of ECs but did not find any stable clusters. Second, transcriptional profiles from EC groups 2, 3, and, partially, group 4 (cyan, green, and red color in the left part of Fig 2) are clearly distinguishable from the healthy control. These groups are similar to each other in terms of transcriptional profiles, but they differ in the magnitude of gene expression changes. For example, EC group 2 was associated with 1063 upregulated genes with log fold change > 0.7 and adjusted p-value < 0.1. However, only 232 of 1063 genes were also upregulated in group 3, and this number increased when the threshold was lowered, e.g., 447 genes were upregulated with log fold change > 0.5 and 559 genes with log fold change > 0.3. Nevertheless, significant numbers of DEGs are unique for each of the three groups, e.g., 315 of 1063 genes are upregulated in EC group 2 but not upregulated in EC group 3 with any log fold change thresholds at p-value < 0.05. Similarly, 84 of 263 genes were upregulated in EC group 3 but not upregulated in EC group 4. The same differences were observed for downregulated genes. Third, EC group 1 (blue color) and a significant portion of cART-treated progressors had transcription profiles opposite those in EC groups 2, 3, and 4 (middle part of Fig 2).

The transcription profiles from ECs and cART-treated progressors cannot be directly compared with untreated progressors’ profiles, since they were measured on different microarray platforms. Instead, we performed a cluster analysis of corresponding log fold changes, calculated by dividing the average expression values in each group to the average expression values in healthy controls (Fig 3).

**Fig 3.**
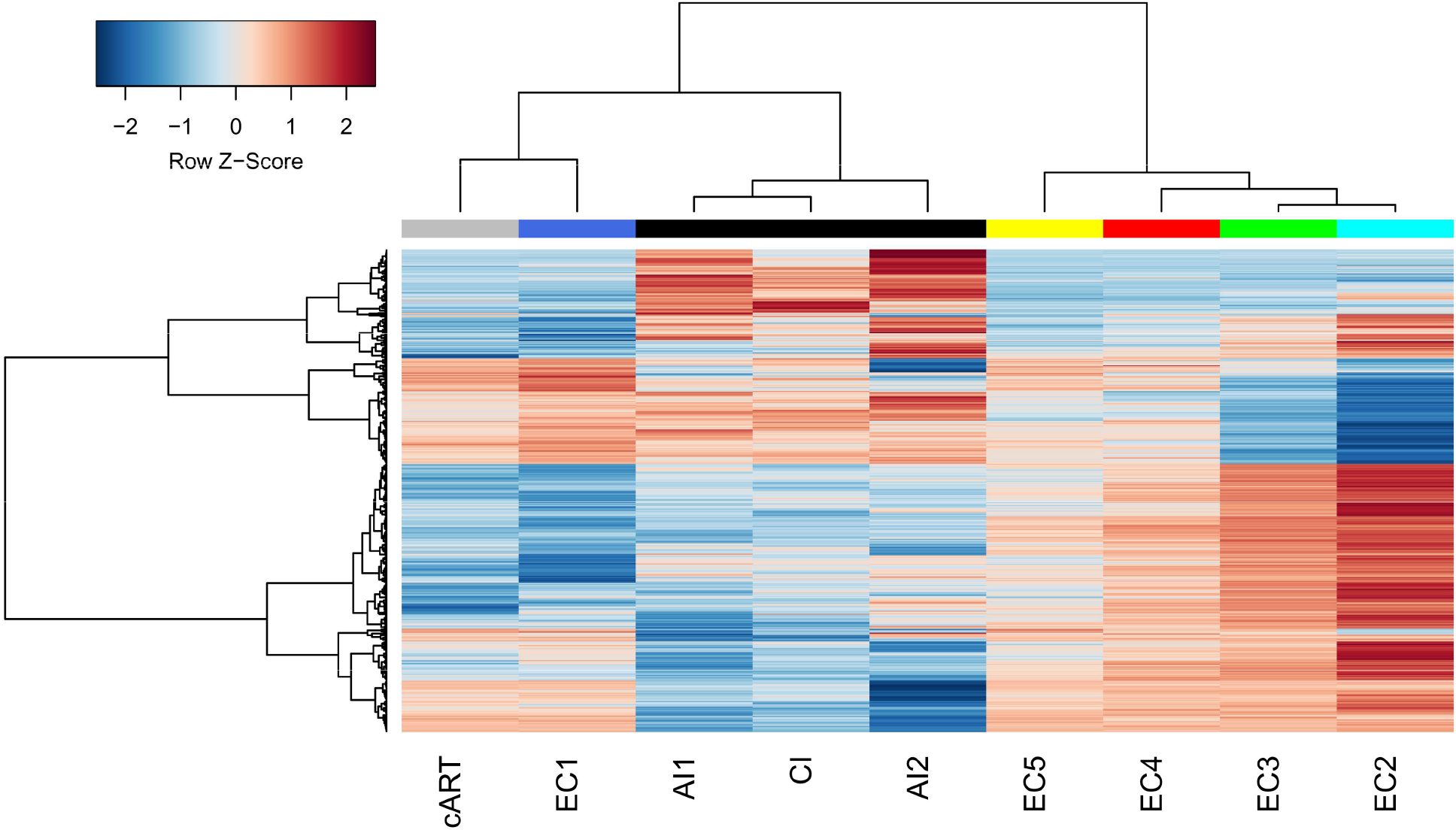
Comparison of log fold changes from five EC groups, cART-treated and untreated progressors. The rows in the heatmap are genes; the columns are groups of HIV-infected individuals. The EC groups 1, 2, 3, 4, 5 (EC1-5) are marked by blue, cyan, green, red, and yellow colors. cART-treated progressors (cART) are marked by grey color. Untreated progressors in acute (AI1 and AI2) and chronic (CI) phases are marked by black color. Only genes, which were differentially expressed in at least one of the investigated groups (Table 2), were used to create the heatmap. Row Z-Score is the number of standard deviations by which the value of log fold change in particular column is above or below the mean value of all groups.

Although transcription profiles from untreated progressors were measured on different platforms, the corresponding fold changes are clustered together to confirm the correctness of the approach used. Profiles from different phases of infection (acute and chronic) are also similar. An earlier study by Hyrcza and colleagues did not reveal DEGs between samples of CD8^+^ lymphocytes from acute and chronic phases (27). Thus, we will further refer to profiles of progressors with no regard to the phase of infection.

Fig 3 shows that EC groups 2 to 5 are clustered together and have transcription profiles different from those of untreated progressors. EC group 1 and cART-treated progressors are clustered together and have transcription profiles distinct from other groups. Thus, three large groups of distinct CD8^+^ cell transcription profiles exist: (1) EC groups 2-5, with a decreasing number of DEGs compared to healthy controls in order from 2 to 5; (2) untreated progressors, whose profiles are different from those of ECs, which may indicate observed dysfunction of CD8^+^ lymphocytes; and EC group 1 and most of the cART-treated progressors, whose profiles are different from healthy controls but opposite to other EC groups and different from untreated progressors.

### Identification of pathways and cellular processes related to identified EC groups, cART-treated and untreated progressors

To identify KEGG pathways (https://www.genome.jp/kegg/pathway.html related to the groups mentioned above, we performed gene set enrichment analysis (see Materials and Methods). We manually checked the positions of DEGs in revealed pathways to filter out nonrelevant ones. For example, if “p53 signaling pathway” was found but the TP53 gene was not differentially expressed, the pathway was removed from further analysis. The obtained list of KEGG pathways is presented in Fig 4.

**Fig 4.**
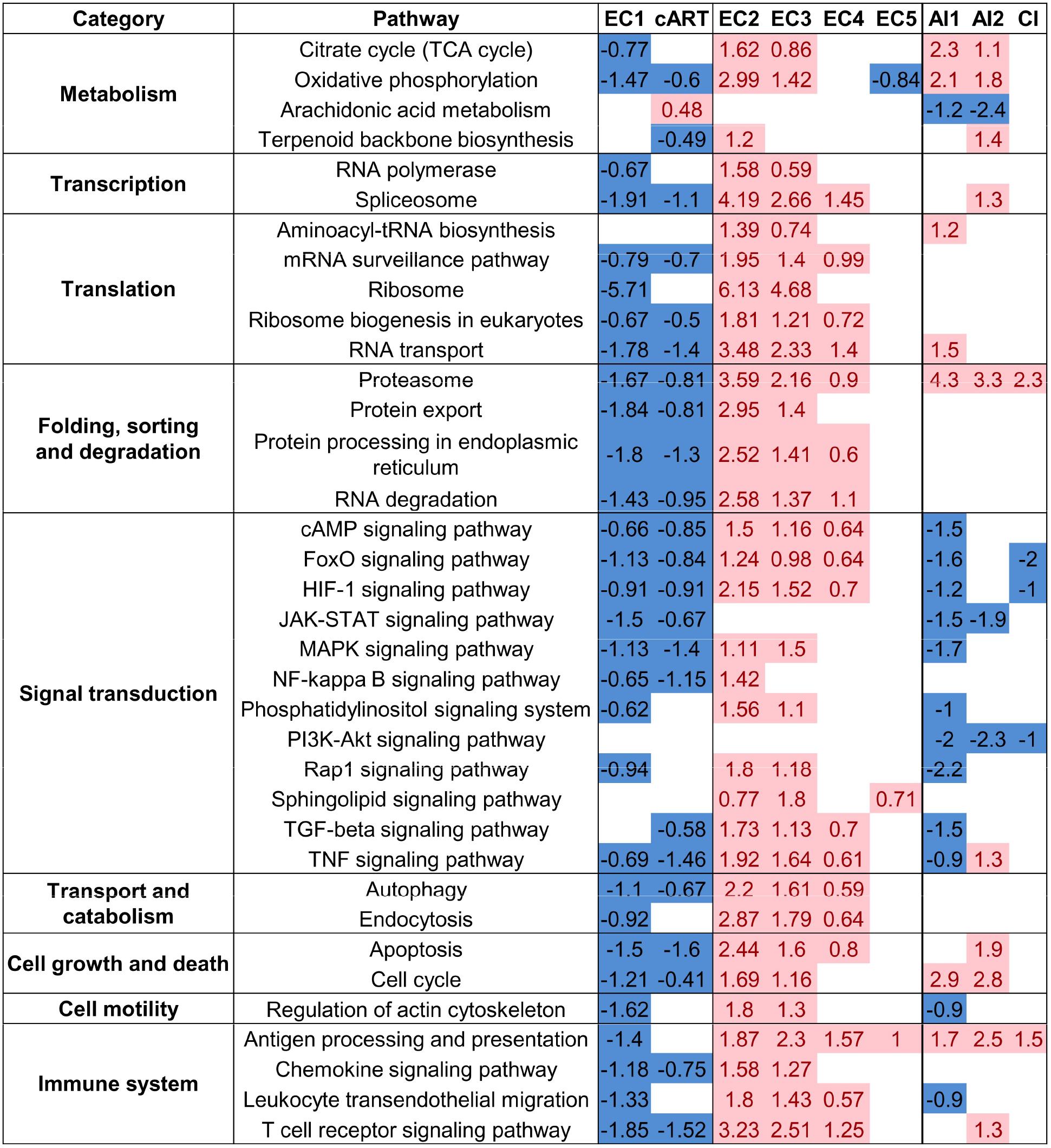
KEGG pathways are differentially regulated in ECs, cART-treated and untreated progressors. The values (columns 3-5) are t-test scores calculated in gene set enrichment analysis (see Materials and Methods). The positive value and red color mean that pathway is up-regulated, whereas the negative value and blue color mean that pathway is down-regulated compared to healthy control. EC 1-5 are groups of ECs; cART is cART-treated progressors, AI1 and AI2 are untreated progressors in the acute phase from two GEO datasets, CI is untreated progressors in the chronic phase.

The key DEGs from pathways related to activation, survival, and immune-specific CD8^+^ lymphocyte functions are presented in Table 3. Here, we considered genes as differentially expressed if the log fold change was more than |0.5| and the unadjusted p-value was less than 0.05. The corresponding thresholds were chosen empirically to balance the number of key genes and differential expression with statistical significance.

**Table 3.**
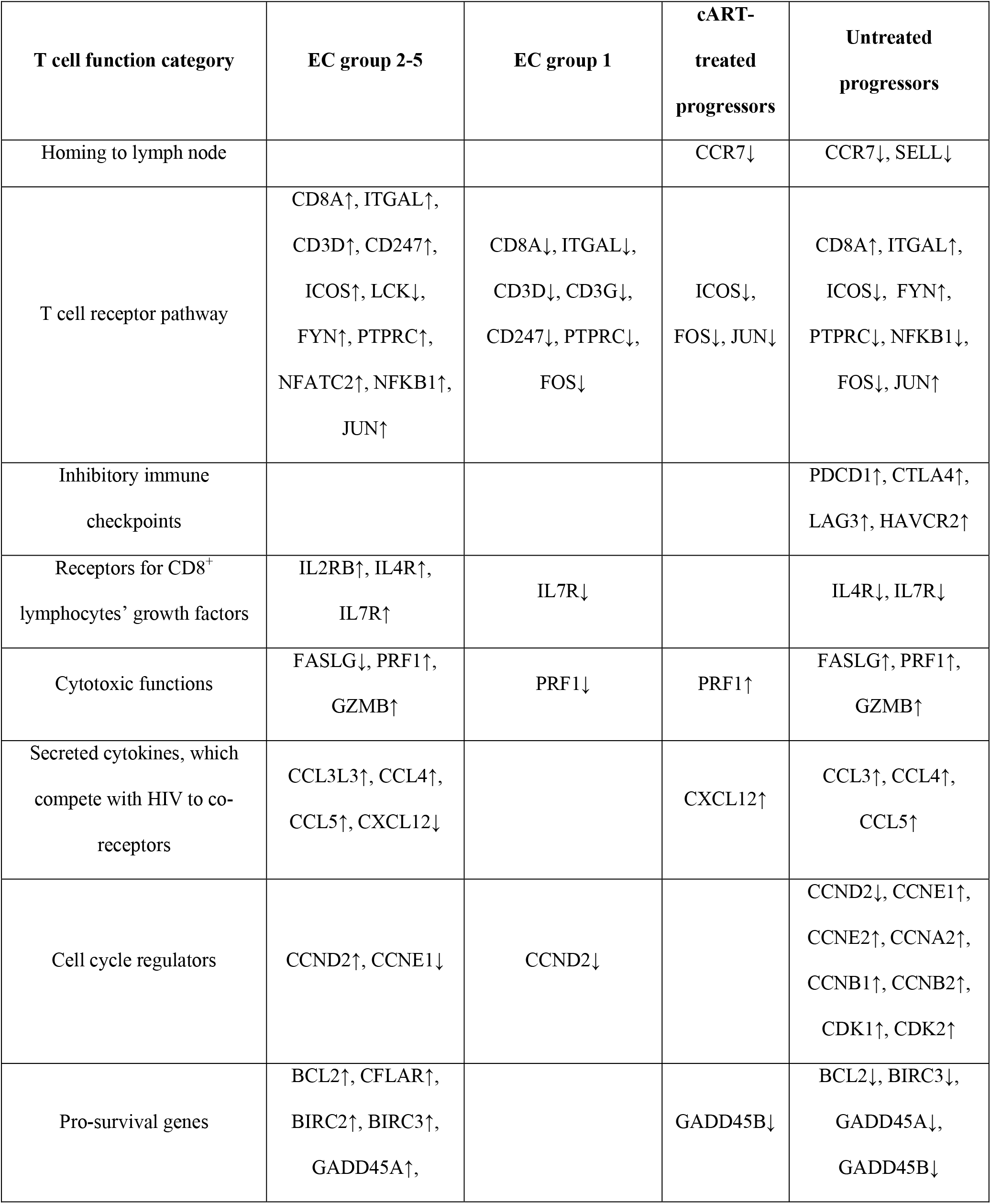
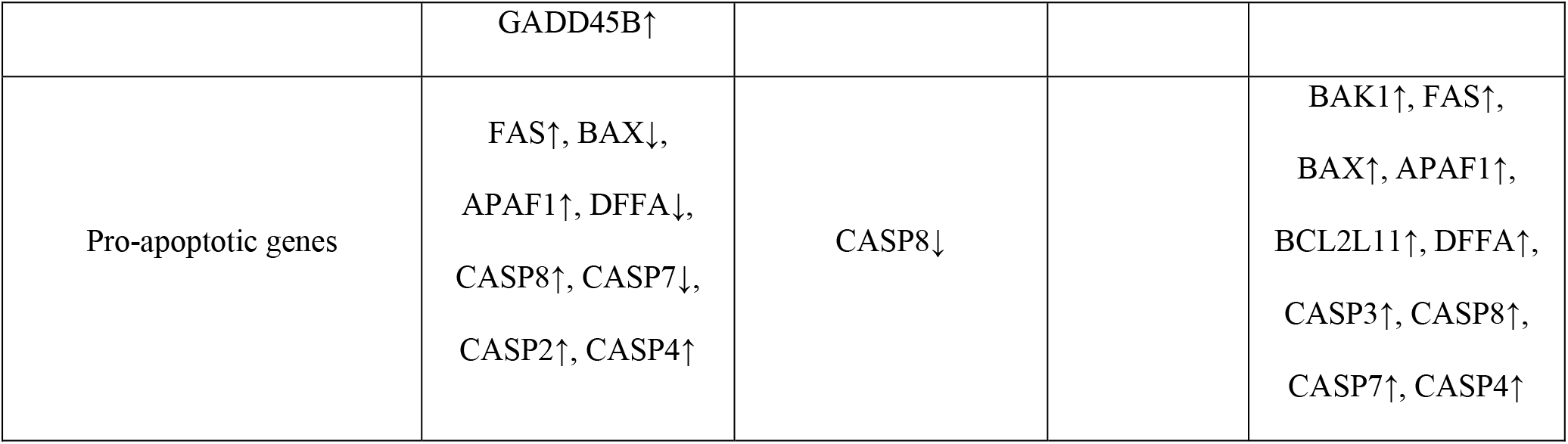
The key differentially expressed genes from revealed pathways representing the most important cellular processes, potentially related to HIV progression.

Fig 4 shows the same relations between EC groups, as well as treated and untreated progressors, as presented in Fig 3. The highest number of pathways was found for ECs. Untreated progressors are associated with fewer pathways, and the direction of regulation of some of them, especially signal transduction pathways, is opposite to most ECs. The direction of pathway regulation in EC group 1 and cART-treated progressors was opposite to EC groups 2-4 at the level of DEGs (Fig 2 and 3).

EC groups 2 and 3 are associated with the same differentially regulated pathways, whereas group 4 is associated with fewer pathways which, however, may be observed because some transcription profiles from EC group 4 are indistinguishable from healthy controls (Fig 2), but other profiles from this group are similar to those from groups 2 and 3. The t-scores obtained for each pathway by gene set enrichment analysis (see Materials and Methods) decreased from group 2 to 4, which indicates the different magnitudes of the pathways’ differential expression. We performed gene set overrepresentation analysis (see Materials and Methods) to find KEGG pathways associated with genes that are differentially expressed in EC group 2 (EC group 3) but not differentially expressed in EC group 3 (EC group 4) (see above). The obtained pathways (Table S2) completely intersected with pathways from Fig 4. Thus, we conclude that EC groups 2 to 4 are similar to each other at the level of pathways but different in the number of DEGs and the magnitude of fold changes. EC group 5 was indistinguishable from healthy controls at the level of pathways.

The most important cellular processes regulated by the identified pathways are as follows: (1) cell metabolism and protein synthesis; (2) T cell activation, migration, and performing CD8^+^ cell-specific functions, including contact cytolysis of target cells and secretion of cytokines; and (3) cell growth, proliferation, and apoptosis. The highest number of revealed pathways was related to cell metabolism and various steps of protein synthesis: from gene transcription to translation, folding, transport, and degradation (Fig 4) (for more details see Discussion).

### Identification of master regulators responsible for the observed transcriptional changes in ECs

Since HIV does not infect CD8^+^ lymphocytes, the observed transcription changes in ECs may be a consequence of the action of cytokines, growth factors, and mediators on corresponding receptors on the surface of cells. We used the Genome Enhancer tool (http://my-genome-enhancer.com) to find MRs, which are the proteins at the top of the signaling network regulating the activity of transcription factors and their complexes and, in turn, are responsible for expression changes observed in ECs (see Materials and Methods). We selected only those MRs whose transcription changed significantly with log fold changes higher than |0.5| and p-values less than 0.5. This was done to filter out irrelevant MRs, which may not influence gene expression in ECs and may not even be expressed in CD8^+^ lymphocytes. The transcription changes of selected MRs themselves mean that they are part of positive feedback loops and are extremely important to maintain the transcription profiles observed in ECs.

We also calculated MRs for cART-treated and untreated progressors for comparison. Most MRs obtained for each group of ECs, cART-treated and untreated progressors are intracellular “hubs,” such as kinases, phosphatases, ubiquitin ligases, GTPases, and transcription factors (Table S3). We focused on receptors because their interaction with corresponding ligands is the first of the consequent events leading to gene transcription changes. As a result of the analysis, we identified 22 receptors, which may be responsible for the observed transcription changes in five EC groups (Fig 5) (for details on receptors, see Discussion). The receptor was selected if the corresponding gene was differentially expressed with a log fold change greater than |0.5| and a p-value less than 0.5 in at least one of five EC groups.

**Fig 5.**
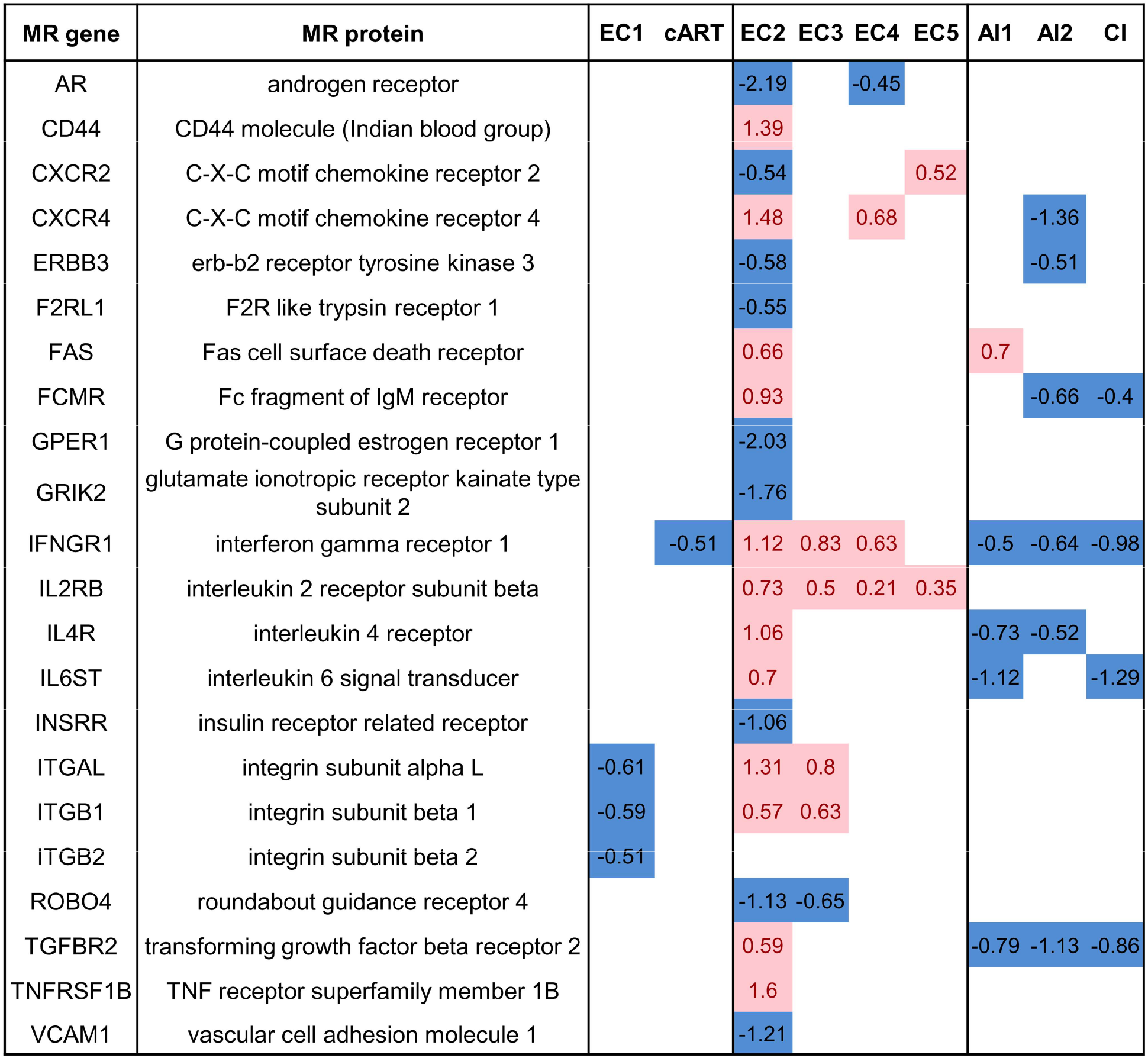
Receptors, identified as MRs, and their transcription changes in EC groups, cART-treated and untreated progressors. EC 1-5 are groups of ECs; cART is cART-treated progressors, AI1 and AI2 are untreated progressors in the acute phase from two GEO datasets, CI is untreated progressors in the chronic phase.

## DISCUSSION

Little is known regarding the mechanisms governing viral control in ECs (11, 13, 18–22, 24), but it appears that cytotoxic and noncytotoxic responses of CD8^+^ lymphocytes play a major role. Naive CD8^+^ T lymphocytes recognize MHC-I restricted HIV antigens presented by dendritic cells in lymph nodes that cause their activation, proliferation, and differentiation into cytotoxic T lymphocytes. After that step, HIV-specific CD8^+^ lymphocytes are capable of recognizing viral antigens in complex with MHC-I on the surface of infected cells and cause their apoptosis by secretion of perforin and granzyme B, as well as by FasL–Fas interaction (11, 13). The virus enters the cell using the CD4 receptor and various coreceptors, including C-X-C chemokine receptor type 4 (CXCR4) and C-C chemokine receptor type 5 (CCR5) (31). Thus, CD8^+^ lymphocytes can also exhibit noncytotoxic anti-HIV functions through the secretion of chemokines that compete with HIV particles for the corresponding coreceptors (19). CD8^+^ lymphocytes from ECs have T-cell receptors capable of broader cross-recognition of mutated epitopes from HIV gag antigens, which may be a consequence of the presence of HLA-I polymorphisms, e.g., HLA-B*57 and B*27, that influence the selection of T cell clones in the thymus (18, 19, 22, 32). CD8^+^ lymphocytes from ECs do not demonstrate an “exhaustion” state and have preserved functions related to cytotoxic and noncytotoxic responses (18, 19, 22, 24), including the ability to degranulate and secrete cytokines with anti-HIV effects, such as MIP-1α/β, RANTES, IFN-γ, TNF-α, and IL-2. They also have a higher proliferation rate and low level of apoptosis compared to progressors.

Total CD8^+^ lymphocytes rather than HIV-specific lymphocytes may be involved in mechanisms of viral control and slow disease progression in ECs. On the one hand, “bystander activation” of non-HIV-specific lymphocytes in ECs does not lead to their dysfunction and apoptosis. On the other hand, non-HIV-specific CD8^+^ cells may still contribute to the control under HIV replication, e.g., by secretion of cytokines that compete with HIV particles for the corresponding coreceptors.

In our study, we analyzed the transcriptional profiles of total CD8^+^ lymphocytes (including HIV- and non-HIV-specific lymphocytes) from ECs, cART-treated, and untreated progressors using the original pipeline. The corresponding analysis allowed us to identify several groups of ECs, which are characterized by distinct transcriptional profiles, up- and downregulated pathways, cellular processes, and MRs. We compared the obtained pathways and MRs to those from cART-treated and untreated progressors to hypothesize the potential mechanisms of slow disease progression.

We identified three large clusters (displayed in Fig 3 and Fig 4) of transcription profiles from the total CD8^+^ cells: (1) profiles from ECs which, in turn, formed four distinct groups (EC groups 2-5); (2) profiles from untreated progressors, which are different from those of ECs; (3) profiles from a small group of ECs (EC group 1) and profiles from cART-treated progressors, which are opposite to other EC groups and different from untreated progressors.

The transcriptional profiles from EC groups 2-5 differ in numbers of DEGs and the magnitude of fold changes (the ratio of the average of expression values in each group to the average of expression values in healthy controls): group 2 is associated with the highest number of DEGs and highest magnitude, whereas group 5 is indistinguishable from the healthy control. Nevertheless, EC groups 2 to 4 were similar at the level of differentially regulated pathways (Fig 4). On the other hand, these groups are different in terms of MRs (Fig 5, Table S3), which regulate the transcription of observed DEGs. EC group 2 was associated with the highest number of differentially expressed MRs and positive feedback loops (see Results), which can explain the highest number of DEGs among all four groups (Table 2). EC groups 3 and 4 are associated with fewer and different MRs. Given the information on DEGs (Table 2), differentially expressed pathways (Fig 4), and MRs (Fig 5, Table S3), we can conclude that the same cellular functions are changed in CD8^+^ lymphocytes from EC groups 2-4 but with different magnitudes, and the initial causes may be different, which is reflected by different MRs.

The identified pathways may indicate preserved functions, survival, and proliferation of CD8^+^ lymphocytes from EC groups 2-4 compared to untreated progressors. Many pathways related to metabolism and protein synthesis were upregulated in EC groups 2-4, whereas fewer corresponding pathways were upregulated in untreated progressors, which may indicate a lower degree of increase in metabolism (Fig 4). The anti-apoptotic genes were upregulated in CD8^+^ lymphocytes from EC groups 2-4 and downregulated in untreated progressors (Table 3). The pro-apoptotic genes were upregulated in untreated progressors, whereas they have a mixed pattern of expression in EC groups 2-4. Thus, CD8^+^ lymphocytes from EC groups 2-4 have a higher ability to survive than untreated progressors; these observations are in accordance with literature data (13, 19). The transcription of cyclins is also dissimilar in EC groups 2-4 and untreated progressors. The only gene coding cyclin D2 is upregulated in ECs, whereas genes coding cyclins E1, E2, B1, B2, and A2 are upregulated in untreated progressors (see Table 3). Since CD8^+^ cells from the blood are asynchronized in cell cycle phases, this observation may indicate cell cycle arrest in the G2 phase. The HIV-1 protein vpr causes G2 arrest in both infected and uninfected cells, whereas the Vif protein promotes the G1 to S phase transition (33, 34). Only cyclins D1 and D2 are expressed in all phases of the cell cycle; thus, the presence of their transcription in EC groups 2-4 and the absence of transcription of other cyclins may indicate normal CD8^+^ lymphocyte proliferation. CD8^+^ lymphocytes from ECs are known to preserve their ability to proliferate compared to progressors (22). The receptors of CD8^+^ cell growth factors were also upregulated in EC groups 2-4 but downregulated in untreated progressors (Table 3). The absence of dysfunction of CD8^+^ lymphocytes in ECs is supported by unchanged transcription of immune checkpoints, whereas the corresponding genes are upregulated in untreated progressors (17) (Table 3). Interestingly, the transcription of genes coding perforin and granzyme B was upregulated in both groups, but the gene coding Fas ligand was downregulated in ECs but upregulated in untreated progressors. CD4^+^ and CD8^+^ T lymphocytes in LTNPs have lower frequencies of apoptosis than progressors, which correlates with a lower frequency of cells expressing Fas and FasL (35).

In contrast, the corresponding pathways associated with cART-treated progressors are differentially regulated in the opposite direction: they are mostly downregulated, whereas the same pathways from EC groups 2-4 are upregulated (Fig 4). This finding may indicate the downregulation of metabolism, activation, growth, proliferation, and other essential processes in CD8+ T lymphocytes from cART-treated progressors. Some corresponding pathways, e.g., JAK-STAT and FoxO pathways, are changed in the same direction in cART-treated and untreated progressors (Fig 4). However, the transcription of many important genes, e.g., pro- and anti-apoptotic genes, is not changed, unlike in untreated progressors (see Table 3). This finding may indicate that many of the considered processes are preserved in CD8+ T lymphocytes from cART-treated compared to untreated progressors, which is in accordance with literature data (24). It is known that low-level viremia (~ 1 copy/mL of plasma) is detectable in most individuals under cART, which leads to residual activation and dysfunction of CD8+ T lymphocytes.

Surprisingly, EC group 1 is associated with transcription changes and pathways similar to those in cART-treated progressors. Thus, EC group 1 may also be associated with some degree of CD8^+^ T lymphocyte dysfunction and may have other viremic control mechanisms that are unrelated to CD8^+^ cells.

The observed transcription changes in CD8^+^ lymphocytes from ECs may be a consequence of the action of various cytokines, growth factors, and mediators on corresponding receptors on the surface of cells. We identified 22 receptors (Fig 5) that were predicted as MRs by the Genome Enhancer tool and may be responsible for the expression changes observed in ECs. Many of the identified receptors are expressed in the opposite direction in ECs and untreated progressors. Taking into account the key roles that these receptors play in gene expression changes in ECs and progressors, they may represent the potential targets of therapeutic intervention, which enables switching of the phenotype of CD8^+^ lymphocytes to those in ECs, decelerating disease progression and potentially increasing the antiviral noncytotoxic response.

Most of the identified receptors are related to various cytokines, which are important for regulating CD8^+^ lymphocyte functions. For instance, the interferon-gamma receptor is a potential MR for DEGs from both ECs and treated and untreated progressors. IFNGR1 gene transcription changed in the opposite direction in ECs and progressors. Interferon-gamma is one of the most important cytokines required to increase CD8^+^ lymphocyte abundance during viral infection and regulate their homeostasis (36, 37). The IL-4 receptor gene is unexpectedly upregulated in EC group 2 and downregulated in untreated progressors. IL-4 is a T helper 2 cytokine that reduces the ability of CD8^+^ lymphocytes to suppress HIV infection (38). Gene IL2RB coding subunit beta of the IL-2 receptor is upregulated in ECs but unchanged in progressors. In contrast to IL-4, IL-2 enhances CD8^+^ T cell anti-HIV activity and optimizes both effector T cell generation and differentiation into memory cells (38; 39). Thus, it appears that the combination of signaling from various receptors, rather than from a single receptor, may cause gene expression changes and functional states observed in ECs and progressors.

Some of the identified receptors, e.g., androgen and G-coupled estrogen receptors, are not typically associated with CD8^+^ lymphocytes; however, they may also contribute to expression changes in ECs. For instance, the androgen receptor gene is downregulated in EC groups 2 and 4, but its expression is not changed in progressors. It was shown that women are significantly overrepresented in the EC population compared to men (22). This finding can be explained by androgens’ influence on the immune system, including CD8^+^ lymphocytes (40). Thus, the androgen receptor may play a significant role in the observed phenomenon.

## MATERIALS AND METHODS

### Transcriptional datasets

The three datasets GSE87620, GSE6740, and GSE25669 with CD8^+^ cell transcriptional profiles from ECs, cART-treated and untreated progressors, and uninfected individuals were obtained from Gene Expression Omnibus (GEO) (https://www.ncbi.nlm.nih.gov/geo). The dataset GSE87620 includes 51 samples from ECs, 32 samples from cART-treated patients, and 10 samples from uninfected people. The GSE6740 and GSE25669 datasets include 5 and 4 samples from untreated progressors as well as 5 and 2 samples from uninfected individuals. The transcriptional profiles were measured on three microarray platforms: Illumina HumanHT-12 V3.0 and V4.0 expression beadchips and Affymetrix Human Genome U133A Array.

### Preprocessing, clustering of samples and identification of differentially expressed genes

The corresponding analysis was performed using various R packages from The Comprehensive R Archive Network (https://cran.r-project.org) and Bioconductor (https://www.bioconductor.org). All steps of analysis were performed separately on each of the three datasets.

Background correction and quantile normalization of transcription data were performed using different functions depending on the microarray platform: function “rma” from the “affy” package for Affymetrix microarray (GSE6740 dataset) and the “neqc” function from the “limma” package for Illumina beadchips (GSE87620 and GSE25669 datasets).

Next, we removed probes that were unexpressed in all samples of the dataset. To filter out corresponding probes on the Affymetrix microarray, we used the “mas5calls” function from the “affy” package and removed probes having an “absent” score across all samples. To do this on Illumina beadchips, we removed probes that have detection p-values greater than 0.05 in all samples of the dataset.

For comparison of DEGs between datasets, we selected only probes having Entrez IDs and only one probe per gene with the highest variance among samples using the “nsFilter” function from the “genefilter” package.

To find potential clusters on heterogenic transcriptional profiles from ECs, we selected 51 corresponding samples and filtered out 50 percent of genes with the lowest variance using the “nsFilter” function. This step was employed to remove genes whose expression is not changed significantly across samples and cannot be useful to find potential EC groups. To find clusters, we used a hierarchical agglomerative clustering approach implemented in the “hclust” basic R function. We choose 1 – Pearson correlation coefficient between pairs of samples as a distance measure, and the “ward.D2” clustering method (41). To estimate the uncertainty in obtained clusters, we performed multiscale bootstrap resampling using the “pvclust” function from the “pvclust” package. For each cluster, “pvclust” calculates p-values, which indicate how strongly the cluster is supported by data: the higher the p-value, the more probable the cluster’s existence. Function “pvclust” provides two types of p-values: the AU (Approximately Unbiased) p-value and BP (Bootstrap Probability) value. The AU p-value is computed by multiscale bootstrap resampling, whereas the BP value is computed by normal bootstrap resampling (42).

To visualize clusters, we used the “heatmap.2” function from the “gplots” R package. This function creates a heatmap with two dendrograms: one for rows, which are genes, and another for columns, which are samples. To create these dendrograms, the abovementioned clustering method (“ward.D2”) and distance measure (1 – Pearson correlation coefficient) were used. The lengths of dendrogram branches are proportional to 1 – Pearson correlation coefficient values. To highlight the differences in gene expression values between samples, row Z-scores were calculated. The row Z-score is the number of standard deviations by which the value of gene expression in a particular sample is above or below the mean value of all samples.

To identify DEGs for each of the obtained clusters of ECs, groups of cART-treated and untreated progressors compared to uninfected people, we used the Linear Models for Microarray Data (Limma) approach (43). The analysis was performed using functions from the “limma” R package. The thresholds on log fold changes and Benjamini-Hochberg corrected (adjusted) p-values were chosen depending on the particular analysis.

### Gene set enrichment analysis

To identify KEGG pathways (https://www.genome.jp/kegg/pathway.html) that were differentially regulated in each group of ECs, as well as in cART-treated and untreated progressors compared to uninfected persons, we performed gene set enrichment analysis (44) using the “gage” function from the “gage” R package. Function “gage” implemented a two-sample t-test for determining the differential expression of genesets, e.g., genes from a particular pathway, between two conditions, e.g., samples from cART-treated progressors and uninfected individuals (45). We selected pathways with an adjusted p-value of less than 0.1, which is the default threshold. The required data on relations between human genes and KEGG pathways were retrieved from the Enrichr database (https://amp.pharm.mssm.edu/Enrichr/#stats).

### Gene set overrepresentation analysis

To identify KEGG pathways associated with genes that are differentially expressed in EC group 2 (EC group 3) but not differentially expressed in EC group 3 (EC group 4), gene set overrepresentation analysis was used (44). This analysis allows the identification of pathways where investigated genes are overrepresented compared to the background gene set, e.g., all genome genes. To perform analysis, we used the function “enrichr” from the “enrichR” R package. We selected pathways with an adjusted p-value less than 0.1, as in the gene set enrichment analysis.

### Identification of master regulators

To identify MRs, we used the Genome Enhancer tool (http://my-genome-enhancer.com) developed by geneXplain^®^ GmbH. Briefly, Genome Enhancer implemented a pipeline including three main steps: (1) analyze promoter regions of genes to predict transcription factor binding sites using positional weight matrices from the TRANSFAC database (46); (2) since it is clear by now that combinations of TFs, rather than a single TF, drive gene transcription and define its specificity, the combinations of TF binding sites called “composite regulatory modules” are identified (47); and (3) reconstruct the signaling pathways that activate these TFs and identify master regulators at the top of such pathways (48, 49). This analysis uses a signaling network from the TRANSPATH database (50).

## ACKNOWLEDGMENTS

This work was supported by Russian Science Foundation Grant (Grant Number 19-75-10097). We are grateful to the geneXplain GmbH for kindly providing access to the Genome Enhancer pipeline.

## SUPPLEMENTAL MATERIAL

**Table S1**. Groups of CD8^+^ lymphocytes’ transcription profiles from ECs.

**Table S2**. KEGG pathways associated with genes, which are differentially expressed in EC group 2 (EC group 3), but not differentially expressed in EC group 3 (EC group 4).

**Table S3**. Master regulators and their transcription changes in EC groups, cART-treated and untreated progressors. The master regulator was selected if the corresponding gene was differentially expressed with log fold changes more than |0.5| and p-value less than 0.5 in at least one of five EC groups. EC 1-5 are groups of ECs; cART is cART-treated progressors, AI1 and AI2 are untreated progressors in the acute phase from two GEO datasets, CI is untreated progressors in the chronic phase.

